# Host trees partially explain the complex bacterial communities of two threatened saproxylic beetles

**DOI:** 10.1101/2024.07.24.604729

**Authors:** Michał Kolasa, Rama Sarvani Krovi, Radosław Plewa, Tomasz Jaworski, Marcin Kadej, Adrian Smolis, Jerzy M. Gutowski, Krzysztof Sućko, Rafał Ruta, Tomasz Olbrycht, Sergey Saluk, Maria Oczkowicz, Łukasz Kajtoch

## Abstract

Microorganisms are integral to ecosystem functioning and host adaptation, yet the understanding of microbiomes in diverse beetle taxa remains limited. We conducted a comprehensive study to investigate the microbial composition of two red flat bark beetle species, *Cucujus haematodes* and *C. cinnaberinus*, and assessed the influence of host taxonomic relatedness and host tree species on their microbiomes. Moreover, we summarize the available data on the microbiome of saproxylic beetles as a reference. We sampled 67 larvae of two *Cucujus* taxa taken from eleven host tree species. 16S rRNA V4 fragment sequencing revealed distinct microbial communities associated with each *Cucujus* species, with host tree species significantly influencing microbiome composition. Alpha and beta diversity metrics indicated significant differences between microbial communities in both, beetle and host tree species. Principal Component Analysis indicated distinct clustering based on host tree species but not for beetle species. This overlap could be attributed to the similar ecology of both *Cucujus* species. The detection of various bacteria, among which some have already been reported in saproxylophagous beetles, suggests that the red flat bark beetles ingest the bacteria via foraging on other wood-dwelling invertebrates. Our findings show the complex interplay between host taxonomy, microhabitat, and microbial composition in *Cucujus*, providing insights into their ecological roles and conservation implications. This research helps to fill the gap in understanding the microbial dynamics of saproxylic beetles, sheds light on factors shaping their microbiomes, and highlights the importance of considering both host species and environmental conditions when studying insect-microbe interactions in forest ecosystems.

## INTRODUCTION

Microorganisms play a pivotal role in the functioning of ecosystems (Van Der Heijden et al., 2008),(Singh et al., 2019). They are not only an element of the environment but also colonize living organisms and often establish dynamic relationships with them [3]. Symbiosis with microorganisms allowed insects to colonize new habitats, explore new food niches, and potentially overcome environmental pressures (Lu et al., 2016). Furthermore, bacterial symbionts can be viewed as an enormous genetic reservoir for insect adaptation to human-induced environmental changes in times of global biodiversity crisis (Feldhaar, 2011).

Some bacteria that live on the surface or inside organisms are neutral for their hosts, whereas others can have negative effects such as diseases, parasitism, decreasing fitness or reproduction of the host (Drew et al., 2021), or have positive effects, like playing a substantial role in provisioning nutrients, acting against pathogens or facilitating development or reproduction (Zug and Hammerstein, 2015). Bacterial taxa associated strictly with their hosts often express some level of co-speciation with the hosts, as their transmission depends on the reproduction of colonized organisms (vertical transmission) (Kikuchi, 2009). However, there is growing evidence that symbiosis, which is considered stable on the evolutionary time scale, undergoes dynamic rearrangement, leading to symbiont replacement (Kolasa et al., 2023, Michalik et al., 2023). On the other hand, the majority of bacteria can easily spread across various individuals and taxa via many horizontal routes of transmission (e.g., common habitats, food, prey, parasites, etc. (Zug and Hammerstein, 2015)). Recently, there has been increasing evidence that horizontal transmission is also important for endosymbionts, enabling their spread but complicating the understanding of host-microorganism relationships (Russell, 2019).

One of the key questions in microbiome analyses is what factors determine its diversity. It can be assumed that the composition of bacteria assemblages would be similar in closely related hosts (species or populations) (Ley et al., 2008). On the other hand, considering the widespread horizontal transfer of bacteria (Russell, 2019), the environment could be expected to determine the microbiome composition in hosts, even if they are not closely related (Wiedenbeck and Cohan, 2011). Many studies have dealt with this issue and examined similarities and differences in microbiome composition or specific bacterial taxa among different hosts, sometimes focusing on taxonomic (phylogenetic) background or various environmental determinants (such as shared habitat, food, or geographic localization of sampled hosts). These results either support the crucial role of evolutionary relations among hosts (Henry et al., 2013) or the importance of shared environmental features, or they demonstrate that both factors contribute to the diversity of examined hosts (Adair and Douglas, 2017). The issue is probably more complex, as many factors influence the diversity of microorganisms observed among hosts in shared (or not) environments. This is still an open question that deserves further studies focusing on some specific systems (host taxa belonging to specific trophic guilds, untypical environments, etc.).

Most studies dealing with the dynamics and functions of bacterial symbionts in insects focus on a few model species, such as *Drosophila* flies (Douglas, 2018), or on some aphids and bees (McLean et al., 2019, Motta and Moran, 2024). However, knowledge of the diversity and roles of symbionts in the vast majority of insects, a group that makes up most of the world’s diversity, is extremely limited (Salem et al., 2023).

Beetles are the most diverse taxonomic order of organisms on Earth inhabiting most terrestrial habitats and occupying nearly all trophic niches (Stork et al., 2015). Hence, there is growing interest in microbiome-focused studies on beetles and their symbionts (Salem et al., 2023). However, most of these studies concentrated on selected beetle taxa, resulting in narrow conclusions. There is a great number of studies that analyze the occurrence and diversity of particular bacteria in beetles, either in selected species or across multiple taxa chosen based on their taxonomic affiliation (e.g., some genera, families), geography (e.g., beetles from a particular area, continent), or environment (e.g., species associated with selected habitats). Only a few studies examined multiple beetle taxa for their entire microbiome (Gurung et al., 2019).

Among beetles, relatively few studies have focused on bacteria of wood-dwelling (saproxylic) taxa, although it is species-rich (Hjältén et al., 2012) and an important group, both from ecological and economic perspectives. It includes many species of conservation concern (threatened species, relicts of primeval forests) or “pests’’ that infest trees and cause damage to timber production (Gurung et al., 2019),. There are several studies on microbiomes of selected families, although mostly restricted to saproxylophages, which are dependent on the wood-digesting bacteria (Rieker et al., 2022). On the contrary, there are missing studies on the microbiome of scavenging or predatory saproxylic beetles, which could be more diverse given the wide range of their diet and their ability to inhabit various tree species.

The genus *Cucujus,* or flat bark beetles, consists of 22 species mainly inhabiting the Oriental zoogeographic region, with several species known from temperate and boreal forests of Eurasia and North America (Horák et al., 2010). The current knowledge of Asian species is poor, as most of these taxa have only recently been described (Hsiao, 2020). On the other hand, European and North American taxa are well known with respect to their biology, ecology, and, since recently, phylogenetics and phylogeography (Kadej et al., 2022, Sikora et al., 2023).

Recent progress in the development of amplicon-based sequencing, coupled with increase in the availability of bioinformatic tools, opened a new avenue for broad-scale studies on microbiome diversity and relationships with its hosts and the environment (Bokulich et al., 2020). However, there are some caveats that we must consider in order to properly identify bacterial communities and understand their impact on hosts (Knight et al., 2018). The lack of negative controls, insufficient decontamination pipelines, and proper quantification methods can all lead to incorrect conclusions. In the present study, we used a customized laboratory and bioinformatic workflow to understand the role of host taxonomy and inhabited tree species in shaping the microbial composition of two flat bark beetle species. Specifically, we verified the following hypotheses:

Hypothesis 1: *Cucujus cinnaberinus* and *C. haematodes* beetles are characterized by a similar core microbiota, indicating their stability over evolutionary time scales.

Hypothesis 2: The microhabitat (i.e., mostly host tree species) mostly determines the microbiome of red flat bark beetles. Beetles, occurring in the same host tree are characterized by the same bacterial clades in high abundance regardless of the species, which is due to horizontal transfer in the shared environment.

## METHODS

### Study species

In Europe, there are only three taxa of *Cucujus* beetles, namely: *C. cinnaberinus* widespread in Central and Eastern Europe (Horák et al., 2010), *C. haematodes* in remnants of primeval forests in Eastern Europe (Kadej et al., 2022), and *C. tulliae* (Bonacci et al., 2012) being endemic to southern Apennines. All species belong to the *C. haematodes* species group but are classified in distantly related lineages (Kadej et al., 2022), (Horák et al., 2010). They mostly occur in forests with substantial amounts and heterogeneity of deadwood – they prefer relatively recently dead trees with a moderate degree of wood decay (Horák et al., 2010, 2011). They are found in various host trees, often in ashes, oaks, or pines, but also in others (Jaworski et al., 2019). It was assumed that these beetles are scavengers or predators (Bonacci et al., 2020). However, recent studies suggest that they likely also consume fungi (Přikryl et al., 2012), although the latter study examined digested remains in the gut, the origin of which was uncertain (e.g., fungi or plant remains could be taken along with animals).

### Sampling

In the present study we used *C. cinnaberinus and C. haematodes* larvae collected from decaying wood of dead trees in Poland and Belarus between 2013-2019. Sampling was limited by the permissions granted by the nature conservation authorities for this project (up to five individuals per site). In total, we collected 30 individuals of *C. haematodes* in six tree species and 40 individuals of *C. cinnaberinus* in eight tree species. Larvae of both *Cucujus* species occurred together in three tree species. Imago and pupae of these beetles were not considered due to difficulties in collection in the field (adults are mostly observed singular and easily disperse over trees, so it is not possible to assign sampling sites to tree hosts; pupae are very hard to find as are hidden deep in wood). Moreover, the goals of this study do not assume to compare the microbiome of various developmental stages of these beetles. Due to the low quality of DNA, three samples were excluded from further analyses, leaving in total 67 samples (28 for *C. haematodes* and 39 for *C. cinnaberinus*) (Supplementary Table 1).

### DNA extraction and amplicon library preparation

DNA was extracted from whole insect larvae using the Nucleospin Tissue kit (Macherey-Nagel), following the manufacturer’s instructions. Samples and negative controls (DNA extraction control, as well as molecular-grade water as a PCR control, Supplementary Table 1) were used for amplicon library preparation following a modified two-step PCR library preparation approach as described by (Glenn et al., 2019). In the first step, the marker 16S rRNA V4 region was amplified using template-specific primers 515F/806R (Apprill et al., 2015; Parada et al., 2016) with Illumina adapter tails. The PCR products were purified using SPRI magnetic beads and used as templates for the second indexing PCR reaction. Pooled libraries were sequenced on an Illumina MiSeq v3 lane (2 × 300 bp reads) at the Institute of Environmental Sciences of Jagiellonian University.

The primer sequences and detailed protocols for amplicon library preparation are listed in Supplementary Table 2.

### Amplicon data analysis

We processed 16S rRNA amplicon data using a custom pipeline based on USEARCH/VSEARCH (available and described in detail at https://github.com/MikeCollasa/Cucujus_project/tree/main). Aware of the limitations of Illumina sequencing, including sequencing errors, chimera formation, and cross-contamination among samples (Kircher et al., 2011), we have invested much effort in developing and optimizing workflows that mitigate these challenges. First, using PEAR, we assembled quality-filtered forward and reverse high-quality reads (Phred score > 30) into contigs (Zhang et al., 2014). Next, contigs were de-replicated (Rognes et al., 2016) and denoised (Edgar, 2016); this was done separately for every library to avoid the loss of information about rare genotypes that could occur during the denoising of the whole sequence set at once (Prodan et al., 2020). The sequences were then screened for chimeras using USEARCH and then classified by taxonomy using the SINTAX algorithm and customized SILVA database (version 138 SSU) (Quast et al., 2013). Finally, the sequences were clustered at a 97% identity level using the UPARSE-OTU algorithm implemented in USEARCH.The product of our custom pipeline were tables with two levels of classification: ASVs (Amplicon Sequencing Variant) (also known as zOTUs - zero-radius Operational Taxonomic Units) describing genotypic diversity and OTUs (Operational Taxonomic Units) – clustering genotypes based on a similarity threshold. ASVs have been used in the past for comparing the microbiota of different bees (Aglagane et al., 2023) or spittlebugs (Kolasa et al., 2023) based on ASVs or 99.5% OTUs. The resulting tables were then used for a series of custom steps. Bacterial 16S rRNA gene data were screened for putative DNA extraction and PCR reagent contaminants using negative controls (blanks) for each laboratory step as a reference. Additionally, based on taxonomic assignment, we used a custom Python script that recognizes and filters out OTUs assigned as mitochondria, chloroplast, Eukaryote, or Archaea. Two specimens with a low number of 16S V4 reads (below 1000) after decontamination (not included in the counts above) were excluded from further analysis.

During decontamination, multiple ASVs were either removed as PCR contaminants or taxonomically assigned as non-bacteria (Supplementary Table 3).

### Diversity Analysis

We used the alpha diversity metric to obtain a single diversity value for each sample, making it suitable for univariate tests, as well as the beta diversity metrics to consider all samples simultaneously, leading to various methods for comparing groups, such as analysis of similarity (ANOSIM) (Clarke, 1993) and permutational multivariate ANOVA (PERMANOVA) (Anderson, 2001).

In this study, several steps were performed using the QIIME2 (Bolyen et al., 2019) tool and R (R Core Team 2020). After obtaining the OTU table, diversity metrics were computed to assess the complexity and richness of the microbial communities. Alpha diversity within individual samples was calculated using observed features (richness), Shannon’s Diversity, Pielou evenness, and Faith’s Phylogenetic Diversity. Beta diversity, which measures dissimilarity between samples, was assessed employing Bray-Curtis and Jaccard distance metrics, Unweighted UniFrac and Weighted UniFrac distances. These distances were visualized using Principal Coordinate Analysis (PCoA) plots.

Kruskal-Wallis one-way ANOVA with Dunn’s post hoc test was used to compare alpha diversities across metadata variables (e.g., Species, Tree, Species_Tree). NCBI BLAST (https://blast.ncbi.nlm.nih.gov/Blast.cgi) was used to investigate ASVs of interest in the data. Reads with unobserved taxa were removed for the entire microbiome. Taxonomic filtering was completed using methods from Callahan et al. (2016). After pruning taxa, samples with less than 1,000 reads were also removed. Taxa were agglomerated by genus before determining relative abundance, leaving 65 samples for beta diversity analyses. Analyses to determine the variance and composition of weighted UniFrac distances grouped by variable were performed using the betadisper and adonis2 functions (PERMANOVA; Anderson, 2001) from the vegan package (v2.6-2; Dixon, 2003). Principal coordinate analysis (PCoA) plots were generated from the UniFrac distances. The plots were generated in Rstudio using the Vegan, Phyloseq, and ggplot2 packages (v3.3.6; Wickham, 2016).

### Differential abundance

For the dataset mentioned above, we performed a differential abundance analysis for identifying microbial taxa that are significantly different between the two beetle species. As per Murdie and Holmes study from 2012 we normalized the count data and performed differential abundance analysis using DESeq2 (Love, 2011) package in association with metagenomeSeq for statistical analysis (Paulson et al., 2013) and phyloseq (Murdie and Holmes, 2013) packages. The rationale behind using DESeq2 is that it utilizes a negative binomial model to account for biological variability. We converted the phyloseq object, which included the OTU table, sample data, taxonomy table, and phylogenetic tree, into a DESeq2 dataset using the (phyloseq_to_Deseq) function and then performed the analysis using the defaults. All tests were corrected for multiple inferences using the Benjamini-Hochberg method to control the False Discovery Rate (Jiang et al., 2017) . We compared Cucujus haematodes to Cucujus cinnaberinus so positive log2 fold changes would indicate higher abundance in Cucujus haematodes as compared to Cucujus cinnaberinus, while negative log2 fold changes would indicate higher abundance in Cucujus cinnaberinus compared to Cucujus haematodes.

## RESULTS

### Microbial composition of *Cucujus* beetles

In 65 samples with at least 1000 reads for the V4 region of the 16S rRNA gene, we compared the presence of more abundant clades, focusing on OTUs that exceeded a relative abundance of 1% in at least one library. We identified 6526 OTUs. Of all the OTUs, 38.4% occurred in both beetle hosts, 31.5% only in *C. cinnaberinus* and 30.1% only in *C. haematodes* (Fig. 1).

**Fig. 1.**
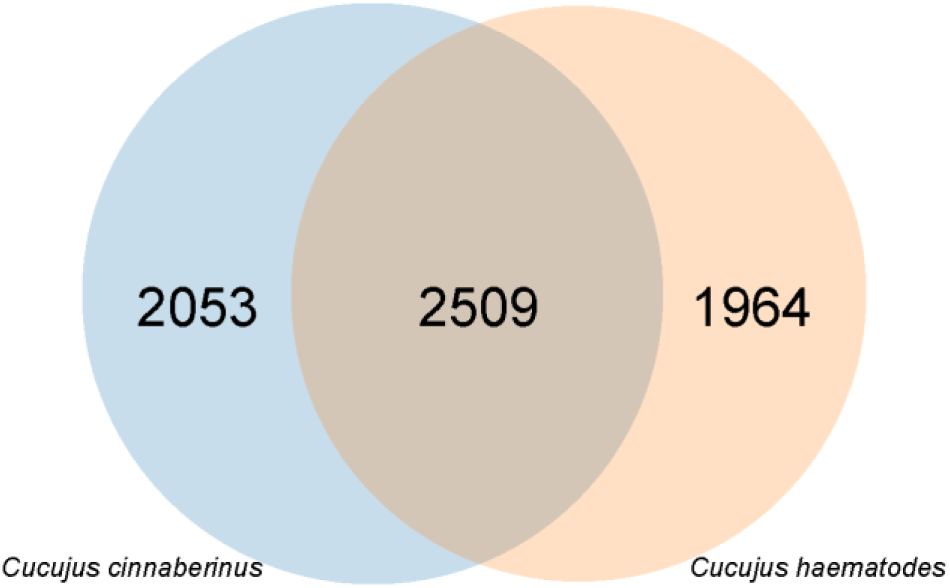
Venn Diagram showing the overlap of bacteria operational taxonomic units found in two studied *Cucujus* beetle species.

The microbial composition of both species showed a diverse community across various taxonomic levels. The phylum level showed a substantially higher diversity with 311 unique classifications, followed by class with 374, order with 970, family with 1484, genus with 2454, and finally species with 1956 unique taxa. The dataset was dominated by Proteobacteria, which accounted for 37.2% of the total taxa identified. Bacteroidota were also predominant (23.9% of the community) followed by Actinobacteriota (9.0%), Verrucomicrobia (7.7%) and Myxococcota (3.4%). Acidobacteria, Planctomycetes, and Firmicutes accounted for 3.0%, 2.1%, and 0.9% of the community, respectively. Notably, there were additional taxa that were not included in the given list, and accounted for a total of 9.0% of the community (Fig. 2).

**Fig. 2.**
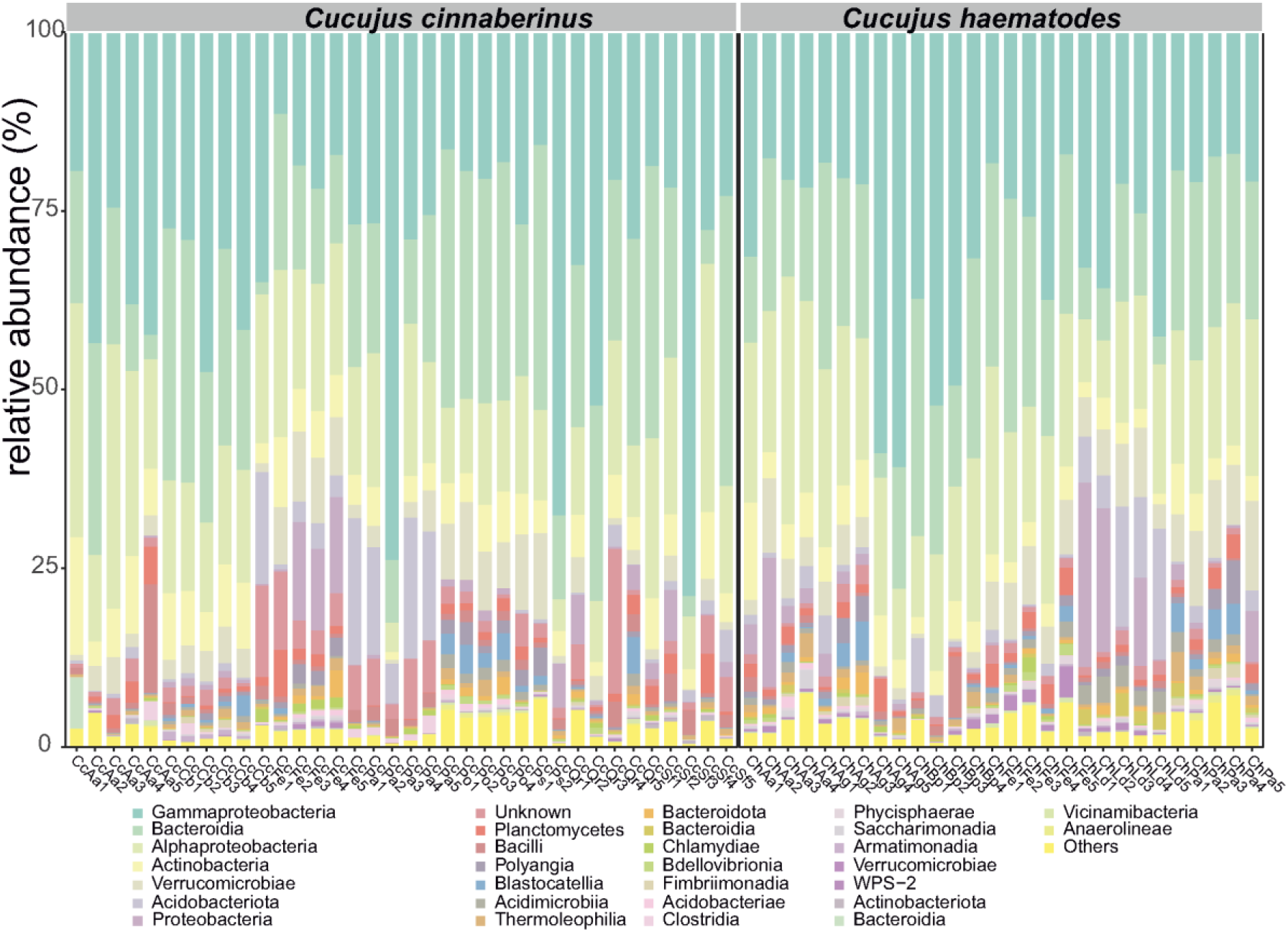
Diagram showing the bacterial classes found in the *Cucujus* beetles examined (shown as relative abundance in each beetle individual examined).

At the family level, the reads for Burkholderiaceae, Nocardiaceae, Chitinophagaceae, Rhodanobacteraceae, Yersiniaceae, Sphingomonadaceae, Pseudomonadaceae, and Weeksellaceae exceed 5% of the total number in the whole dataset. At the genus level, the following bacteria predominate: *Rahnella/Serratia, Pseudomonas, Alkanindiges, Arachidicoccus*, and *Burkholderia-Caballeronia-Paraburkholderia*, followed by *Variovorax, Dyella, Rhodanobacter,* and others (Supplementary Fig. 1).

### Differential abundance analysis

A differential abundance analysis between the species *Cucujus haematodes* and *Cucujus cinnaberinus* showed a total of 99 operational taxonomic units (OTUs) which passed the significance for p-value adjust < 0.05 (Supplementary table 4, Supplementary Fig. 2). Notably, the log2 fold change (MLE) for OTUs revealed substantial differences between the two species, with some OTUs exhibiting markedly higher abundance in *C. haematodes* when compared to *C. cinnaberinus* (Supplementary Fig. 3, 4). For example, among most abundant genera, *Rahnella* displayed significantly low abundance in *C. haematodes*, while*Burkholderia-Caballeronia-Paraburkholderia* or *Dyella* demonstrated a considerable overdominance in *C. haematode*. Among less abundant bacteria a very high overdominance in *C. haematodes* was found in case of Fimbriimonadaceae and Legionellaceae (log2FoldChange 26.18 and 25.47, respectively), whereas high overdominance in *C. cinnaberinus* was detected for Xanthomonadaceae and Spirosomaceae (log2FoldChange -25.76 and -27.15, respectively). Additionally, the adjusted p-values indicated statistical significance for a majority of the comparisons, emphasizing the robustness of the results (Supplementary table 6). We also identified a higher proportion of OTUs with high abundance (53%) compared to low abundant ones (46%), suggesting potential species-specific microbiome (Supplementary Fig. 3, 4) in both *C. haematodes* and *C. cinnaberinus*.

### Alpha diversity

The Kruskal-Wallis test calculated for four metrics (OTUs, Shannon, Faith, and Pielou) measured between beetle species showed significant differences only for OTUs. Significant differences were found between the host trees for Shannon, Faith, and Pielou indices, whereas all metrics showed significant differences between beetles and host trees (Table 2).

**Table 1.** Sampling design of the two *Cucujus* beetles from selected host trees (values indicate numbers of *Cucujus* larvae collected from particular trees).

| *Cucujus* species | Host tree |  |  |  |  |  |  |  |  |  |  |
| --- | --- | --- | --- | --- | --- | --- | --- | --- | --- | --- | --- |
|  | *Betula pendula* | *Fraxinus excelsior* | *Alnus glutinosa* | *Carpinus betulus* | *Populus* sp. | *Quercus robur* | *Salix fragilis* | *Larix decidua* | *Abies alba* | *Picea abies* | *Pinus sylvestris* |
| *C. haematodes* | 4 | 5 | 5 | - | - | - | - | 5 | 4 | 5 | - |
| *C. cinnaberinus* | - | 5 | - | 5 | 4 | 5 | 5 | - | 5 | 5 | 5 |

**Table 2.** Results of the statistical comparison of the microbiome alpha diversity metrics (Kruskal-Wallis test) and the beta diversity metrics (Permutational multivariate analysis of variance, PERMANOVA).

| Kruskal-Wallis test | OTUs |  | Shannon |  | Faith |  | Pielou |  |
| --- | --- | --- | --- | --- | --- | --- | --- | --- |
|  | *H* | *p* value | *H* | *p* value | *H* | *p* value | *H* | *p* value |
| Species | 4.391 | 0.0361 | 2.505 | 0.113 | 2.304 | 0.129 | 1.131 | 0.287 |
| Tree | 16.343 | 0.090 | 19.544 | 0.033 | 21.952 | 0.015 | 22.895 | 0.011 |
| Species Tree | 37.317 | 0.0003 | 32.79 | 0.001 | 39.159 | 0.0001 | 32.629 | 0.001 |
| PERMANOVA | Bray–Curtis |  | Jaccard |  | Weighted UniFrac |  | Unweighted UniFrac |  |
|  | pseudo-*F* | *p* value | Pseudo-*F* | *p* value | Pseudo-*F* | *p* value | Pseudo-*F* | *p* value |
| Species | 2.721 | 0.001 | 1.951 | 0.001 | 2.505 | 0.013 | 2.386 | 0.001 |
| Tree | 2.969 | 0.001 | 2.146 | 0.001 | 2.868 | 0.001 | 2.297 | 0.001 |
| Species_Tree | 3.567 | 0.001 | 2.413 | 0.001 | 3.565 | 0.001 | 2.848 | 0.001 |

### Beta diversity

PERMANOVA calculated for four metrics (Bray-Curtis, Jaccard, Weighted Unifrac and Unweighted Unifrac) proved to be significant for all comparisons (between beetle species, between host trees and between beetles and host trees) (Table 2).

Adonis was used to calculate the F.Model value for various factors. The high R2 value of 5.40 emphasizes that the model explains a significant 58.4% of the total variation in microbial community composition when considering both “Species” and “Tree”. Looking at the individual factors, the “Tree” factor accounts for 41.3% of the total variation, while the “Species” factor explains 4.6% (Supplementary Table 5).

### Principal Component Analysis

PCA calculated using Bray-Curtis dissimilarity matrix showed that there is no clear separation of beetle individuals belonging to two studied *Cucujus* taxa (Fig. 3), although beetles tended to split along the principal component 2-axis which explained only a small part of the total variance (9.2%). When grouping individuals on a PCA plot according to their collection in host trees, there were some visible clusters with the most pronounced samples from poplars, larches or birches, whereas the remaining host trees did not form separate clusters (f. 5). This also applies to trees that are shared by both beetles (ashes, pines and firs), as individuals of *C. cinnaberinus* from these trees do not overlap in the PCA plot with individuals of *C. haematodes* found in the same host tree (Fig. 3).

**Fig. 3.**
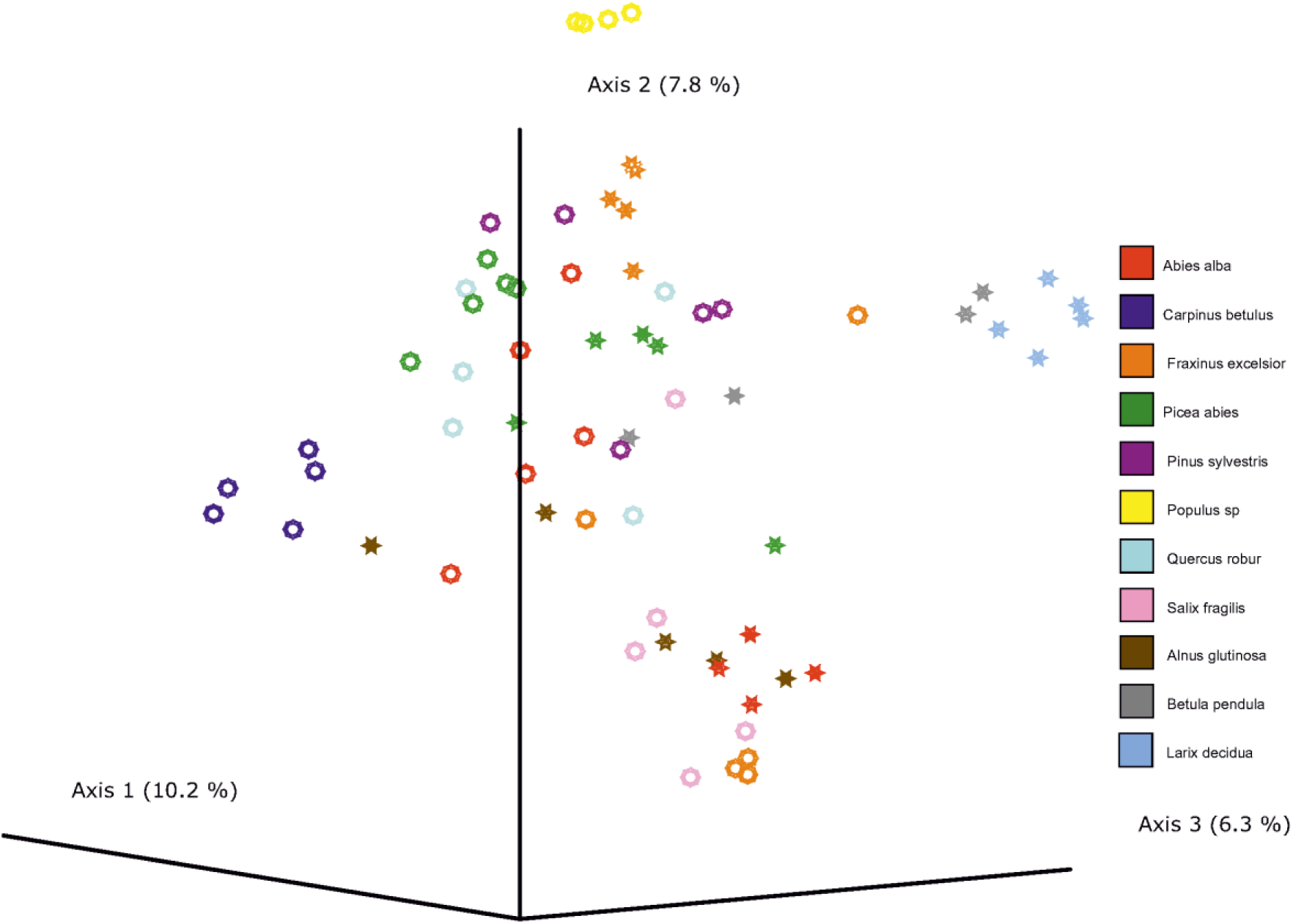
The Principle Component Analysis plot showing the position of examined *Cucujus* beetles based on the dissimilarity of their microbiomes and grouping by host trees in which beetles were collected. Circles – *C. cinnaberinus*, stars - *C. haematodes*.

## DISCUSSION

### Bacteria inhabiting *Cucujus* beetles

Saproxylic beetles are known to host various bacterial taxa, enabling them to synthesize nutrients in deficient environments (Ziganshina et al., 2018). Although our results indicate that *Cucujus* beetles are characterized by a relatively numerous microbiome suggestive of complex interaction with the surrounding environment, we lack clear evidence for its function. Proteobacteria, Bacteroidota, and Actinomycetota dominate the microbial community in red flat bark beetles. These bacteria tend to be ubiquitous in humid environments such as soils, peatlands, and freshwaters, and many of their members have also been reported from forest ecosystems (Humbert et al., 2009). The dominance of such taxa is unsurprising taking into account the microhabitat occupied by red flat bark beetles – dead trees in moderate stages of decay. These beetles are usually found under the loose bark of trees dead for two to four years and prefer to live on moist trunks (with “dark” wood infested with fungi) (Horák et al., 2010, 2011). This microhabitat is shared with many saproxylophagous organisms, including larvae of bark, longhorn, and jewel beetles (Grove, 2002). Although *Cucujus* larvae were considered scavengers or predators that feed on invertebrates (Bonacci et al., 2020), recent studies indicate that they also have fungi and wood in their gut (Přikryl et al., 2012). Consequently, the microbiome composition is of mixed origin, probably at least partially coming from the prey. Many of the bacteria found in *Cucujus* beetles were also reported in other saproxylic beetles, which supports the assumption that red flat-bark beetles share a significant part of their microbiome with their prey (Kajtoch et al., 2019).

Some of the most abundant genera found in *Cucujus* beetles have also been described in other beetles. *Rhanella*/*Serratia* has been reported from the gut of bark beetles (*Dencroctonu*s spp. and *Ips* spp.) and several species of longhorn beetles (Cerambycidae) (Hernández-García et al., 2017). The *Caballeronia* (formerly *Burkholderia*) clade is shared by *Cucujus* spp. and *Bolitophagus reticulatus* (Tenebrionidae); *Dendroctonus* spp. (Curculionidae); and *Monochamus* spp. (Cerambycidae) (Kaczmarczyk-Ziemba et al., 2019, Guo et al., 2020). *Dyella* is known from *Prionoplus reticularis* (Cerambycidae) (Reid et al., 2011). The abovementioned bacteria are ubiquitous in humid environments, including forests (Jeon et al., 2021). In contrast, *Alkanindiges, Arachidicoccus*, and *Rhodanobacter*, have not been reported to dominate in saproxylic beetles other than *Cucujus*, but were described as a part of rhizosphere or soil component (Mehlferber et al., 2021). *Pseudomonas*, which is frequent in the microbiome of *Cucujus*, has also been reported from jewel beetles (*Agrilus planipennis*), bark beetles (*Dendroctonus* spp., *Ips typographus*), and long-horn beetles (*Monochamus* spp., and *Stromatium barbatum*) (Peral-Aranega et al., 2020). Recently, bacteria of the genus *Pseudomonas* have been proposed as nutritional symbionts associated with bark beetles (Falqueto et al., 2022). However, without a more solid molecular approach combined with experiments, we cannot determine the role they may play in the biology of red flat bark beetles. Noticeable, relatively few bacteria found in red flat bark beetles are known to be responsible for the nutrient supply or degradation of substrates such as cellulose (lignin) or chitin, as is the case with other beetles that feed directly on wood or fungi (Shelomi and Chen, 2020). This finding confirms previous findings indicating that *Cucujus* beetles are not saproxylophagous – they do not eat wood or fungi that live on decayed wood and are likely scavengers or predators of invertebrates (Bonacci et al., 2020). Our findings can shed new light and open up new avenues for studies focusing on microbiome dynamics in wood-dwelling beetles, as there are virtually no prior studies describing microbiomes of saproxylic beetles not being saproxylophagous. The only other study on the microbiome of scavenger/predatory saproxylic beetle was work conducted on *Ampedus pomorum* (Elateridae) (Samoilova et al., 2016), in which specimens were mainly infected with Actinobacteria and γ-Proteobacteria with *Streptomyces* predominating. It seems that scavenging or predatory beetles have very different bacterial communities, which probably depend on the diet of particular beetle species or even individuals. Larvae of scavenging/predatory beetles are much more mobile than sedentary larvae of saproxylophagous taxa, as they have to actively search for food (unlike saproxylophages, which forage in their surrounding habitat), so they have many more occasions to come into contact with various microbes.

### Diversity of bacterial communities among and between red flat bark beetles

Our results indicate that bacterial communities of *Cucujus* beetles are abundant and diverse. However, to fully describe their microbiome, an analysis of their structure is required. Our results indicate that although the microbiome of *C. cinnaberinus* is significantly different from that of *C. haematodes*, there is no obvious pattern that distinguishes the beetles of these two species from each other. At the same time, there is a high level of dissimilarity of particular OTUs found in these two beetle hosts. Among many bacteria OTUs showing a different abundance in these beetle hosts, two dominant are known also from other saproxylic beetles: Rahnella was the dominant genus at various stages of Dendroctonus spp. (Curculionidae) and Dyella in Prionoplus reticularis (Cerambycidae) (Reid et al. 2011, Pineda-Mendoza et al. 2022).

Abovelisted findings contradict the hypothesis that the bacteria were inherited from the common ancestor of the two *Cucujus* species. Such a coevolution scenario would be indeed implausible, considering the distant phylogenetic relationships between these taxa, although both belong to the *C. haematodes* species group (Kadej et al., 2022). If the hypothesis is to be positively verified, this might require the study of additional species of flat bark beetles or a targeted study of specific bacteria, as shown in the case of *Monochamus* (a saproxylic longhorn beetle) by Kajtoch et al. (2019). Instead of attributing similarities in the microbiome of *Cucujus* beetles to vertical transmission via common ancestors, we investigated the influence of host trees as a crucial factor for microbiome diversity in saproxylic beetles.

We hypothesize that beetles sharing a common microhabitat, particularly the deadwood of certain tree species, would show a greater bacteria exchange between different taxa. However, our analyses only partially support this hypothesis, as no clear pattern emerged. While there is evidence, where we see some clustering of certain samples based on OTUs in the beta diversity plots, that some individuals found in certain tree species have diverse bacterial assemblages, the overall picture remains nuanced. This suggests that several factors besides the microhabitat influence the composition of the beetles’ microbiome.

In certain tree species, such as poplars, birches, and larches, bacterial communities show greater similarity when collected from beetles inhabiting the same host. In contrast, the microbiomes of different beetle species collected from the same trees, such as ashes, pines, and firs, do not show clustering based on the specific host tree. These results and the fact that some are shared between *Cucujus* beetles and other saproxylic taxa (mainly saproxylophages or mycophages), strongly suggest that flat bark beetles are scavengers and/or predators, and their rich bacterial assemblages originate from diverse preys eaten by the larvae. It is plausible that certain bacteria found in *Cucujus* beetles, especially those typically of environmental origin, were acquired from the moist environment of dead tree wood where the larvae live. Similarly, bacteria associated with fungi or those that compete with fungi could likely be attributed to the movement of *Cucujus* larvae under the bark of dead trees, commonly inhabited by detritivorous fungi (Parisi et al., 2018). However, to fully understand the interplay between fungi and bacteria, further analyses are needed that incorporate the diversity of fungal taxa.

## CONCLUSIONS

The techniques used for microbiome characterization come with numerous caveats that are only sometimes properly considered or implemented (Knight et al., 2018). Being aware of these caveats, we have implemented our customized workflow accompanied by the use of negative controls on every laboratory step, from DNA extraction to PCRs and library preparation and bioinformatic pipeline of high-throughput amplicon data (Kolasa et al., 2023). By implementing spike-in-based quantification, we demonstrated that *Cucujus* beetles are characterized by a rich microbiome that is challenging to interpret. This finding is in line with reports on insects (Kolasa and Łukasik, 2023), and there are cases where insects’ microbiomes lack structure (Coon et al., 2014). In a host infested by ubiquitous bacteria, it is unlikely that significant structuring would be observed. Only when symbionts contribute to the fitness of the host (e.g., by provisioning nutrients, defense against pathogens or parasites, and playing a significant role in proper development), patterns in their prevalence are observed (Gupta and Nair, 2020).

## ACKNOWLEDGEMENTS

We want to thank Monika Prus-Frankowska for helping process quantification samples in the laboratory.

The part of this study (sampling and DNA isolation) was financed by the grant of the National Science Center Poland (UMO-2021/43/B/NZ9/00991, PI – Ł.K.).

## ETHICAL STATEMENT

The sampling required permits in Poland: DLP-III-4102-265/ 21003/15/MD, DOPozgiz-4200/ I-124/ 3303/10/JRO, DZP-WG. 6401.01.20.2018. eb, DZP-WG. 6401.01.30.2018.eb.1.

## Supplementary files

**Supplementary Fig. 1.**
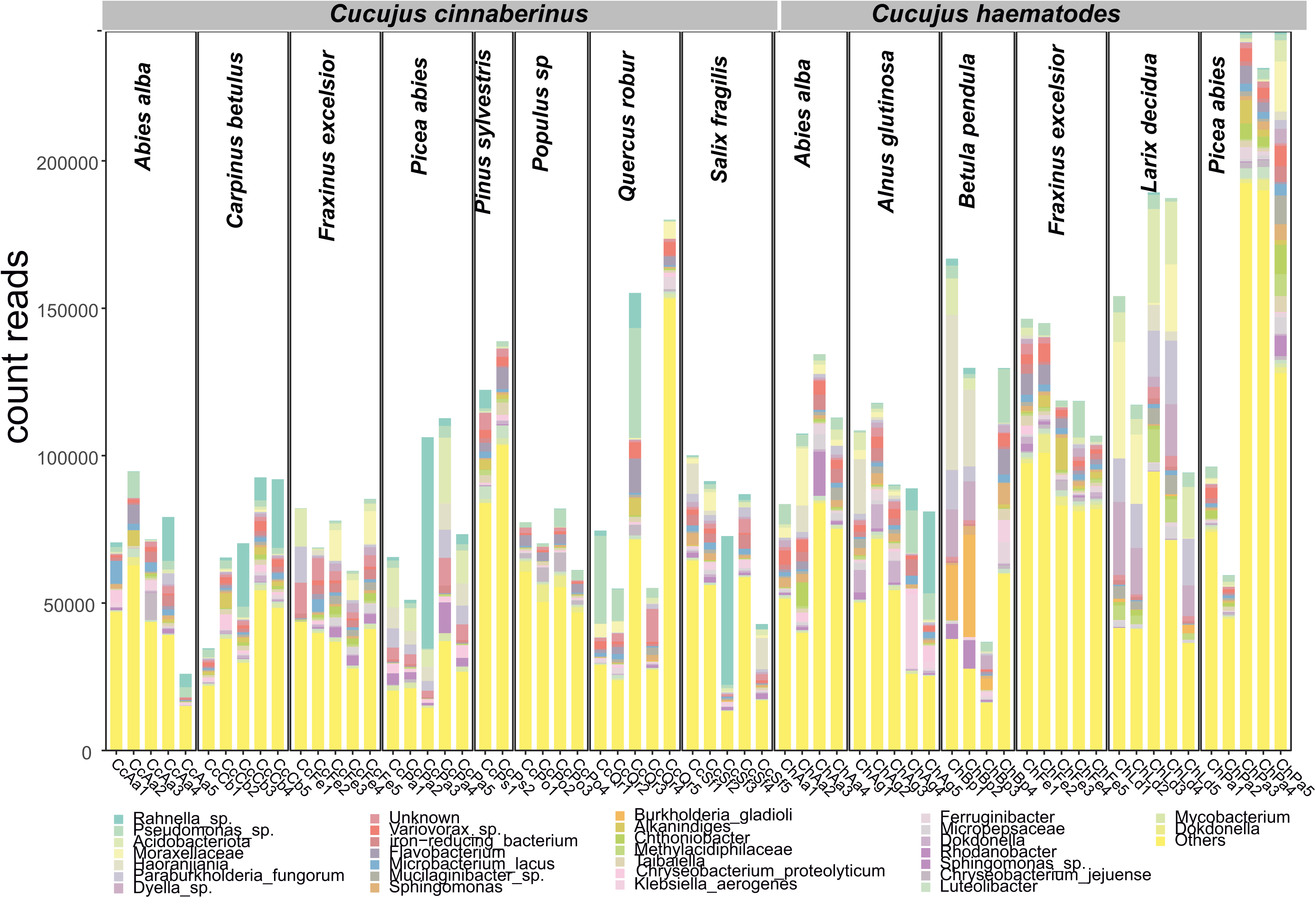
Abundance plot showing the top 30 most abundant bacterial taxonomic units for all examined *Cucujus* beetles.

**Supplementary Fig. 2.**
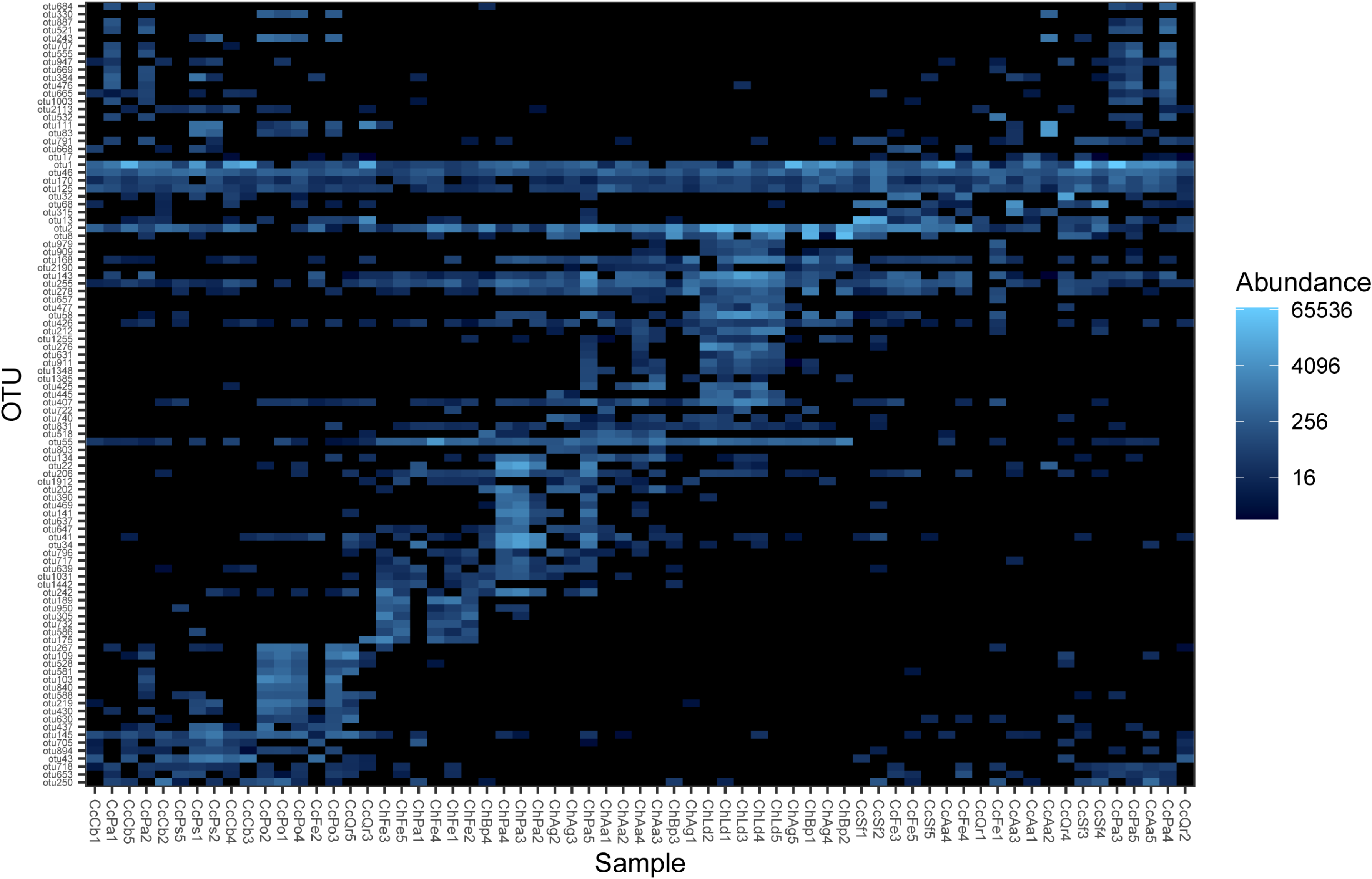
Heatmap of abundance of bacterial operational taxonomic units (OTU) in examined Cucujus beetles.

**Supplementary fig. 3.**
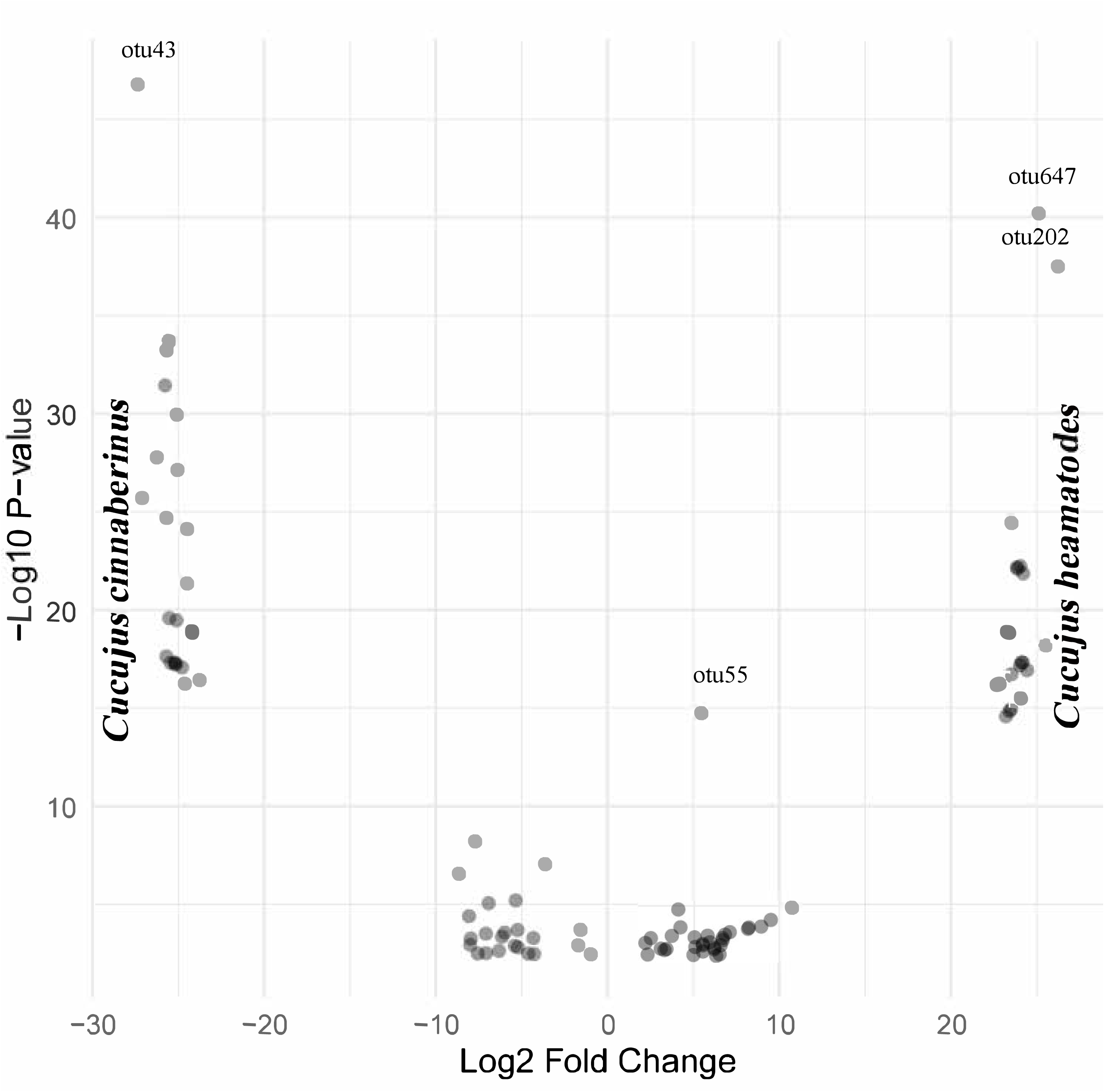
Volcano plot showing operational taxonomic units significantly differentially abundant in examined Cucujus beetles. X-Axis: Positive values indicate OTUs that are more abundant in *Cucujus haematodes*, while negative values more abundant in *C. cinnaberinus*. Y-Axis: Higher values on this axis indicate more statistically significant changes.

**Supplementary fig. 4.**
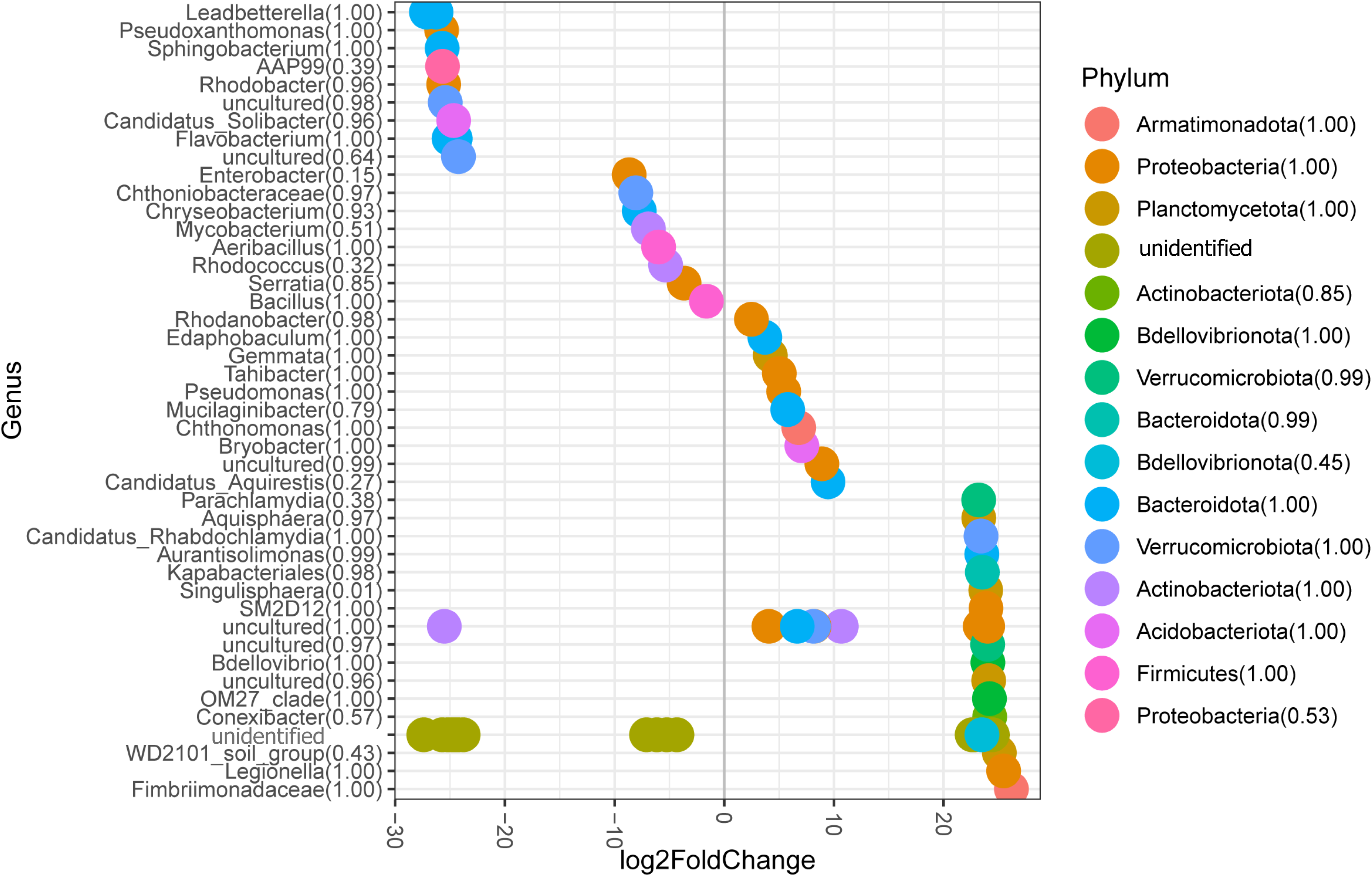
DotPlot showing differential abundance of bacteria genera by phylum detected in examined Cucujus beetles. This dotplot shows the log2 fold change of genera, colored by their respective phyla, indicating their differential abundance. The genera are ordered by their maximum log2 fold change values, from highest to lowest, within each phylum category. Phylum Order: The genera within each phylum are sorted by the maximum log2 fold change observed, providing a hierarchical view of the most significantly changed genera within each phylum. Genus Order: Within the plot, the genera are also sorted by their log2 fold change values, allowing for easy identification of the most differentially abundant genera. The x-axis represents the log2 fold change, with a vertical gray line at zero to distinguish between genera that are more abundant (positive values) versus those that are less abundant (negative values). The y-axis lists the genera, organized by their log2 fold change within each phylum.

**Supplementary Fig. 5.**
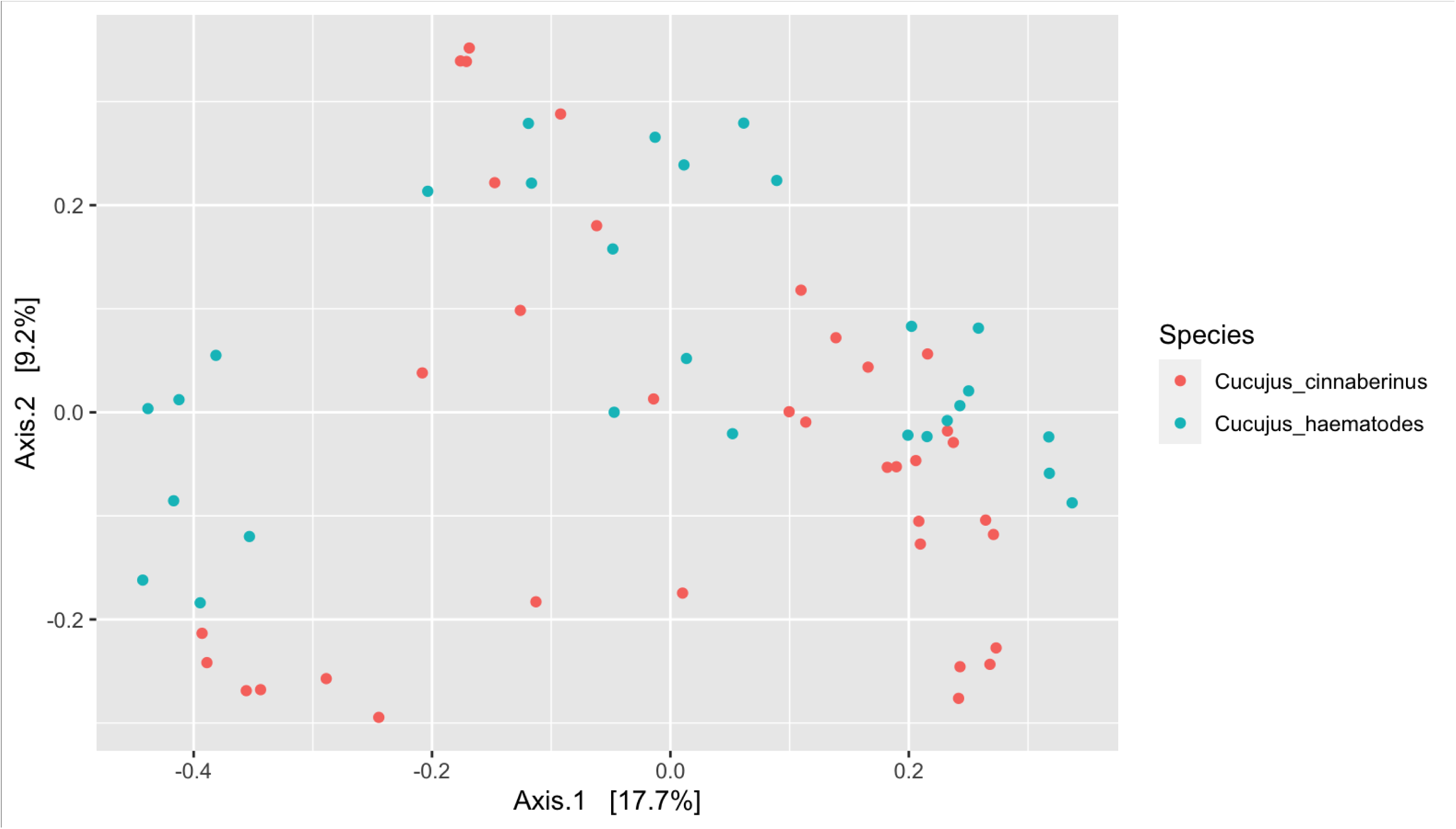
Principal Component Analysis calculated using microbiome dissimilarity (Bray-Curtis) between two examined *Cucujus* beetles.

**Supplementary table 1.**
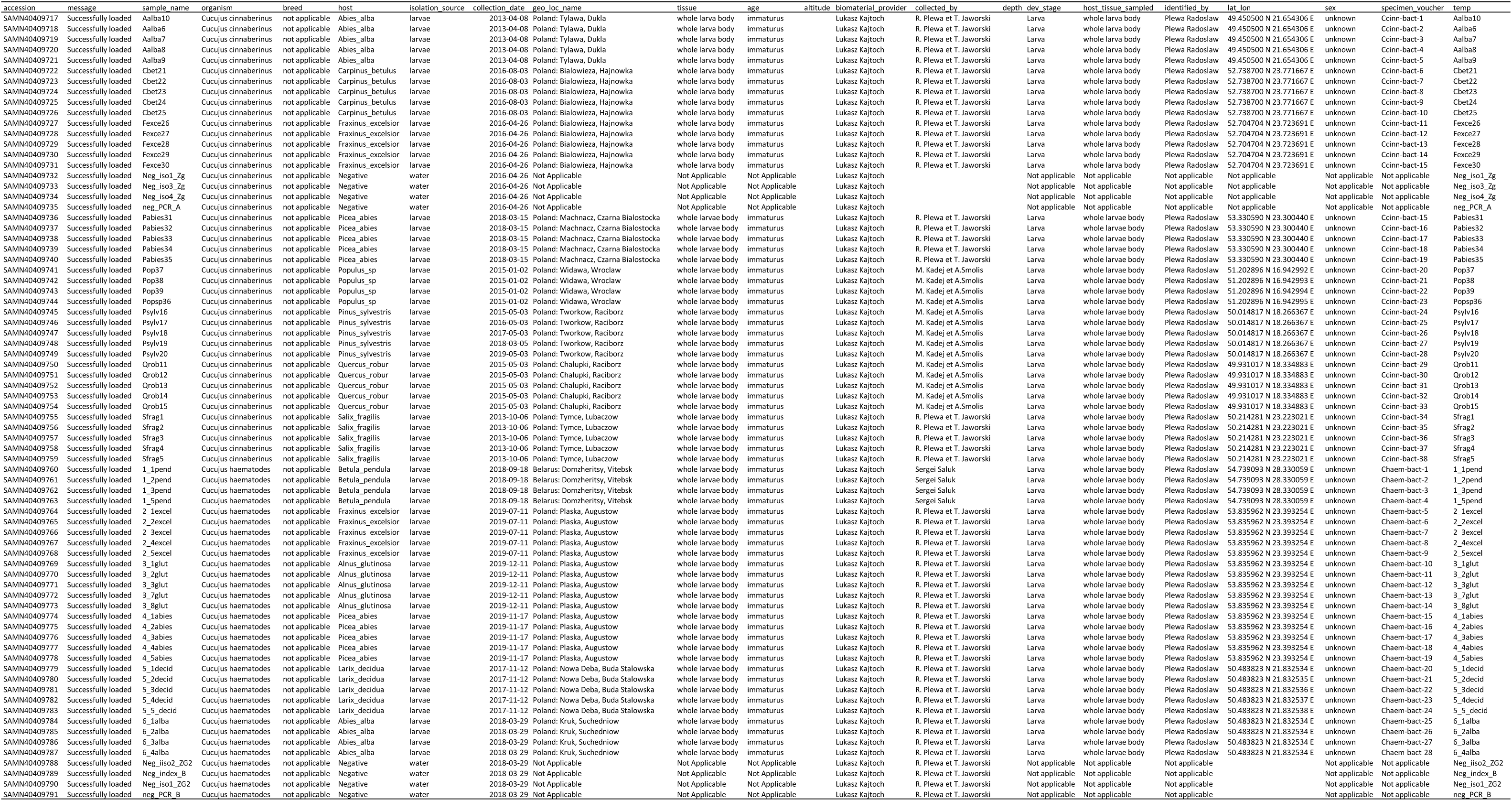
Attributes of *Cucujus* beetle samples used for microbiome sequencing.

**Supplementary table 2.**
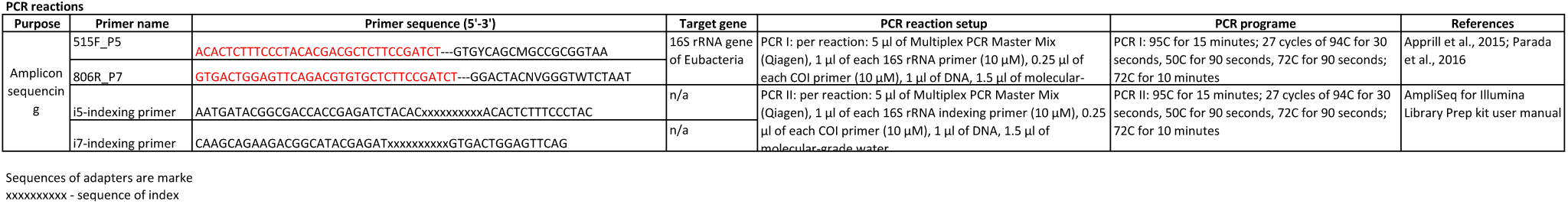
Primers and protocols used for microbiome sequencing.

---

**Supplementary table 3.**
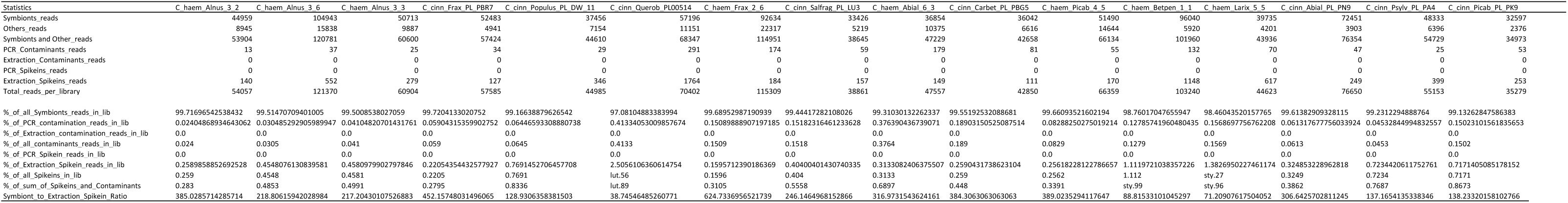
One of the output tables from the custom decontamination with the information about decontamination level and spike-in:symbiont ratio required to calculate absolute abundance.

**Supplementary table 4.**
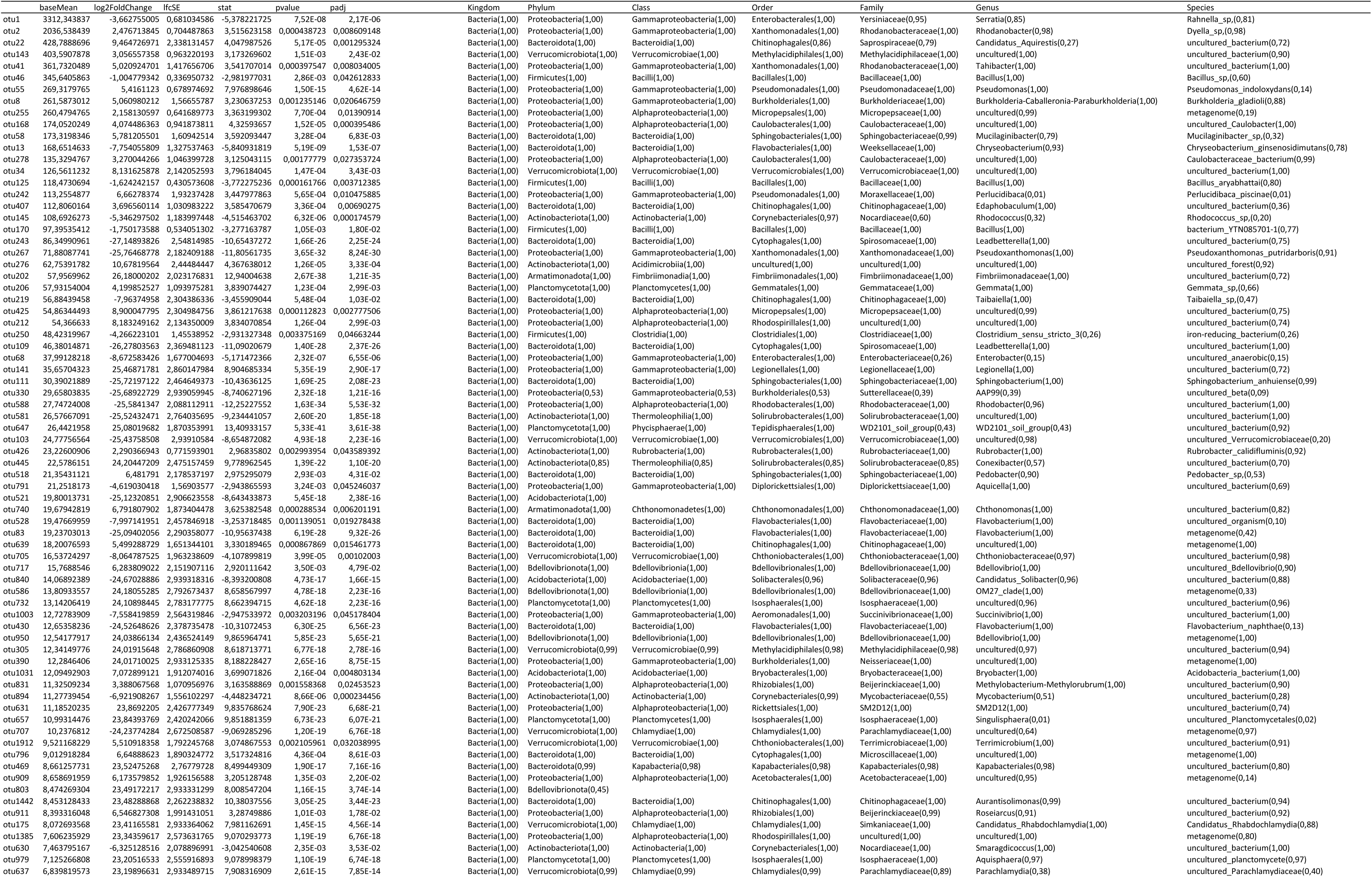
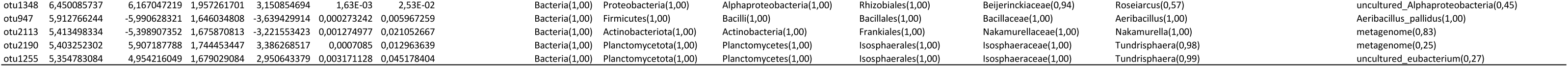
Results of differential expression analysis implemented on operational taxonomic units (otu) determined in two saproxylic beetles: *Cucujus haematodes* and *C. cinnaberinus* . Taxonomic assigment of selected otus is provided in the right part of the table.

**Supplementary table 5.**
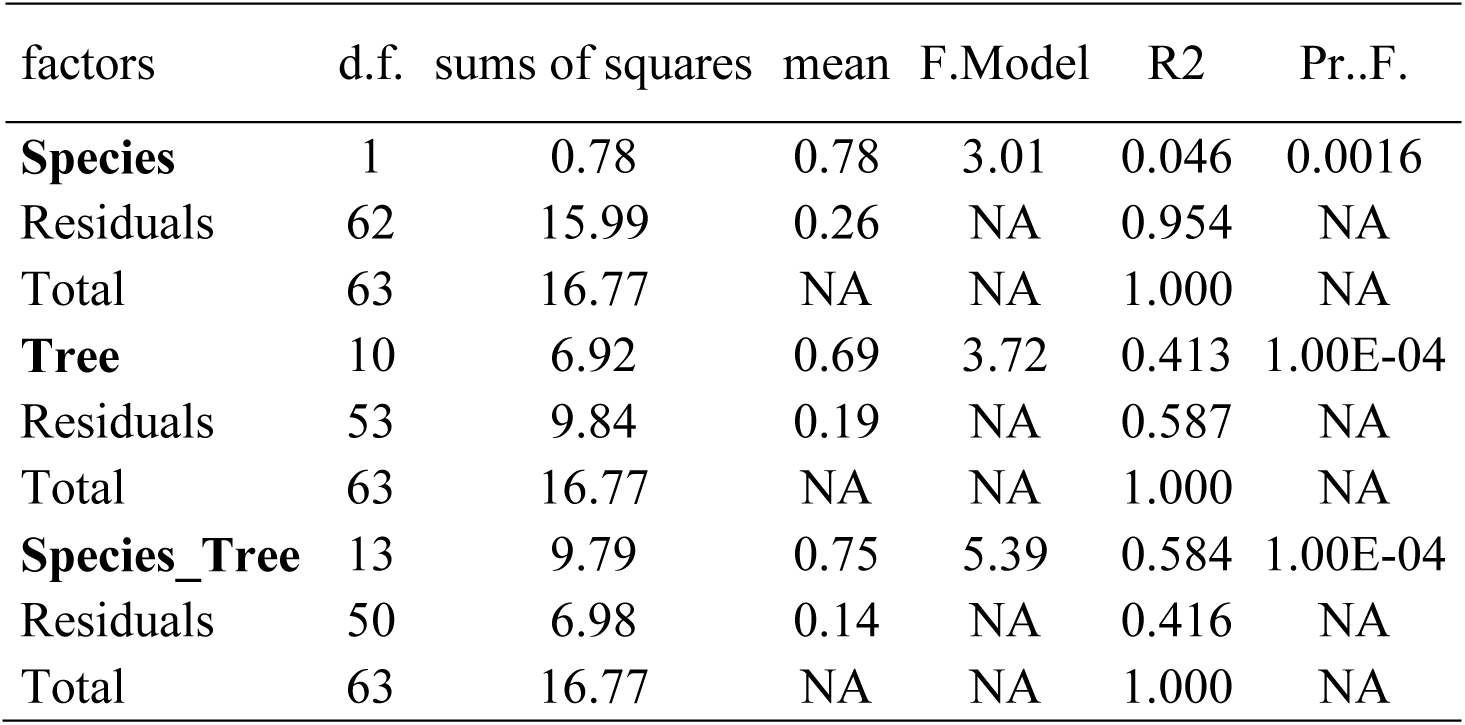
Permutational Multivariate Analysis of Variance calculated on three levels: species (beetle), tree (host tree) and species_tree (beetle species x host tree).

